# Isobaric labeling update in MaxQuant

**DOI:** 10.1101/2024.09.30.615788

**Authors:** Daniela Ferretti, Pelagia Kyriakidou, Jinqiu Xiao, Shamil Urazbakhtin, Carlo De Nart, Jürgen Cox

## Abstract

We present an update of the MaxQuant software for isobaric labeling data and evaluate its performance on benchmark datasets. Impurity correction factors can be applied to labels mixing C- and N-type reporter ions, such as TMT Pro. Application to a single-cell multi-species mixture benchmark shows high accuracy of the impurity-corrected results. TMT data recorded with FAIMS separation can be analyzed directly in MaxQuant without splitting the raw data into separate files per FAIMS voltage. Weighted median normalization, is applied to several datasets, including large-scale human body atlas data. In the benchmark datasets the weighted median normalization either removes or strongly reduces the batch effects between different TMT plexes and results in clustering by biology. In datasets including reference channels, we find that weighted median normalization performs as well or better when the reference channels are ignored and only the sample channel intensities are used, suggesting that the measurement of reference channels is unnecessary when using weighted median normalization in MaxQuant. We demonstrate that MaxQuant including the weighted median normalization performs well on multi-notch MS3 data, as well as on phosphorylation data. MaxQuant is freely available for any purpose and can be downloaded from https://www.maxquant.org/.

## Introduction

Isobaric labeling^1–3^ is a genuine way to achieve high levels of multiplexing combined with high precision for relative quantification in shotgun proteomics experiments. The MaxQuant^4,5^ software supports the analysis of isobaric labeling data since longer and here we present an update of the isobaric labeling capabilities and showcase their performance on benchmark data with known quantitative ratios as well as on relevant biological datasets. In particular we find that the PSM-level weighted median (WM) normalization^6^ introduced earlier is very effective at removing or reducing the batch effects caused by experimental designs whose samples span more than one n-plex of isobaric labels. A well-established strategy for quantitative comparisons across isobaric labeling batches is to include a reference channel, containing the same sample in each n-plex, which has ideally a large overlap in proteins with all the samples to be measured, for instance by pooling all samples. Despite its obvious benefits, it comes at the cost of losing some channels for the actual samples and potentially not being able to quantify some proteins when they are missing or low abundant in the reference channel. Therefore, here we perform a cost-benefit assessment of reference channels on relevant isobaric labeling datasets in combination with the WM normalization in MaxQuant. Furthermore, we demonstrate the application of the latest version of MaxQuant to datasets enriched for post-translational modifications, such as phosphorylation.

## Experimental section

### Datasets

*Dataset 1*^7^ is a mixed species single cell proteomics dataset which was created to investigate the influence of the carrier channel in single cell proteomics by mass spectrometry^8^ (SCoPE-MS). It was labeled with TMTpro^9^ 16-plex reagents. The channel 126 was the carrier proteome channel constructed with the mixture of Homo sapiens (human), Escherichia coli (E. coli) and Saccharomyces cerevisiae (yeast) proteomes, it has the ratio of 14, 42, 98, 210 and 434 to the single cell proteome channel. The channel 127 C was left empty, For each single cell proteome channel there were 200 pg of proteins, in channel 127N and 130N the mixing ratio of proteins from human: E. coli: yeast was 2:1:1. Samples were measured on Orbitrap Exploris 480 mass spectrometer coupled to the Evosep One system as described in ref^7^. The raw data were downloaded from the ProteomeXchange Consortium^10,11^ under the identifier PXD027742. The fasta files of the reference proteome including isoforms from UniProt^12^ (Release-2023_05, UP000000625: 4415 proteins, UP000002311: 6090 proteins, UP000005640: 104473 proteins) were used for database search. Proteome Discoverer 2.4 and MaxQuant version 2.6.3 were used for this dataset.

*Dataset 2*^13^ is a large-scale dataset comprising a quantitative proteome map of the human body. Samples originate from the Genotype-Tissue Expression (GTEx) project containing samples from 54 tissues of 948 post-mortem donors^14,15^. 201 GTEx samples from 32 different tissue types of 14 normal individuals were studied by proteomics which covered all major organs. TMT 10-plex was used in a MultiNotch MS3 mode^16^, with the tissue samples randomized so that each plex consisted of an assortment of tissues and a reference sample in two channels. To increase the proteome coverage, each TMT 10-plex sample was extensively fractionated (12 fractions/sample). The Waters online nano 2D LC system was coupled to an Orbitrap Fusion. We downloaded the raw data from the PRIDE repository with the identifier PXD016999. MaxQuant version 2.4.9 was used to process the raw files. The fractions were combined in MaxQuant by specifying the same ‘Experiment’ name for each of the 12 fractions belonging to a sample. The search was performed against the human UniProt database (release version 2023_03, canonical + isoforms sequences) containing 103,789 protein entries.

*Dataset 3*^17^ is coming from a double-proteome proteomics experiment that includes sub-datasets generated from both MS2 and synchronous precursor selection^16,18^ (SPS)-MS3 acquisition strategies, each recorded using both TMT10/11-plex and TMTpro 16-plex reagents. The measurements were conducted on Orbitrap Fusion Tribrid mass spectrometer. The experiment involved five samples containing a constant 50 μg of HeLa digest and varying amounts of yeast digest (1, 1.75, 3, 5.2, and 9 μg). Each sample was analysed with six replicates, resulting in 30 samples per labeling design. In the 11-plex dataset, three separate 11-plex experiments were set up, each consisting of two replicates for each of the five different ratio samples, plus a reference sample containing 50 μg of HeLa digest and 3.9 μg of yeast digest. In the 16-plex Dataset, two 16-plex experiments were prepared, each consisting of three replicates for each of the five different ratio samples, plus a reference sample identical to the one used in the 11-plex dataset. Mass spectrometry proteomics data (Thermo raw files) were obtained from the ProteomeXchange Consortium under the identifier PXD014750. The raw files were searched using MaxQuant (version 2.6.3) against yeast and human databases downloaded from the UniProt Knowledgebase (release version 2023_03, canonical + isoforms sequences). The FASTA files contained 103,789 protein entries for human and 6,090 protein entries for yeast.

*Dataset 4*^19^ is a time series dataset with nine time points that was recorded to compare MS3 SPS to MS2 high field asymmetric waveform ion mobility spectrometry^20–22^ (FAIMS) based TMT quantification. HEK293 cells were exposed to heat stress for up to 9 hours in 1-hour increments and labeled with different TMT10 reagents. Triplicate injections were performed for the SPS method while three different CV combinations were used with FAIMS. Samples were analyzed on an Orbitrap Fusion Tribrid. Raw data with the dataset identifier PXD009547 were downloaded from the MassIVE repository. The human UniProt database used in dataset 2 and dataset 3 was used also in this case. MaxQuant version 2.6.3 was used.

*Dataset 5*^23^ comes from a phospho-enriched, two-proteome multiplex (TMTpro 16-plex) proteomics experiment that investigated reporter ion interference and ratio compression. The experiment used proteomes from human Jurkat E6-1 cells and the yeast strain Saccharomyces cerevisiae (W303-1A). Human peptides, which made up the majority of each sample, were used as a stable quantitative background, while yeast peptides were varied in defined human:yeast ratios of 0:0, 100:0, 100:6, 100:9, and 100:12. These defined ratios allowed for fold change calculation and differential expression testing. Each experimental condition was prepared in triplicate, resulting in a total of 12 peptide samples, the channel 126C was left empty. The experiment was performed using a Q Exactive HF-X Orbitrap mass spectrometer. Mass spectrometry proteomics data (Thermo raw files) were obtained from the ProteomeXchange Consortium under the identifier PXD040449. The raw files were searched using MaxQuant (version 2.6.3) against yeast and human databases downloaded from the UniProt Knowledgebase (release version 2024_09, canonical + isoforms sequences). The FASTA files contained 104,949 protein entries for human and 6,091 protein entries for yeast.

### Processing of isobaric labeling data in MaxQuant

All quantification is based on the reporter ion signals in MS2 or MS3 spectra. For that purpose, MaxQuant collects for each protein group all PSMs (peptide-spectrum matches) of all associated identified peptides according to the specified protein group-level and PSM-level FDRs, which are by default both 1%. Peptides that occur in more than one protein group are by default assigned to one of them according to the razor peptide principle. Isobaric matching between runs^24^ can add MS2 spectra that were not identified with stringent FDR criteria, but that are located on three-dimensional MS1 features obtained by matching between runs. Further MS2-level labeling specific filters can be applied, as for instance, based on the precursor ion fraction or base peak ratio, but these were not used here, except for dataset 5. Several output files, such as peptides.txt, modificationSpecificPeptides.txt, sites.txt, evidence.txt and msms.txt, contain columns named ‘Reporter intensity …’ which are filled with reporter intensity signals of a single PSM in msms.txt and the sum of reporter ion intensities over all applicable PSMs for all other tables. An exception is the proteinGroups.txt file, which contains the summed reporter intensities only when the parameter ‘Normalization’ is set to ‘None’, and otherwise the accordingly normalized values. The tables in proteinGroups.txt, peptides.txt, modificationSpecificPeptides.txt and sites.txt contain separate sets of reporter intensity columns for each experiment in the experimental design representing a data matrix over all samples that can be conveniently uploaded in downstream data analysis software, such as Perseus^25^.

### Isotope impurity correction

Let y_1_,…,y_n_ be the uncorrected reporter intensities within one n-plex as they are measured and x_1_,…,x_n_ the as yet unknown corrected intensities. A is an nxn matrix whose rows are mixing weights normalized to 1, as they can be obtained from the purchased batch. These are related by

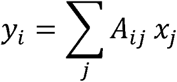

which is a homogeneous linear equation that needs to be solved for x. To improve the stability of the solution, we remove all index values that have zero entries in y from y, x and rows and columns of A, obtaining a generically solvable linear system of equations with a unique solution for x, which we solve by LU decomposition. Negative elements in the solution for x will be set to zero. The values in x are reported as ‘Reporter intensity corrected’ in the output tables, with everything else being the same as for the ‘Reporter intensity’ columns.

### Weighted median normalization

Previously, we introduced weighted median normalization^24^ for protein group level quantification of isobaric labeling data. The main idea is to combine the reporter intensities in a channel over multiple PSMs by a weighted median of ratios of sample channel intensities to a reference intensity. The reference intensity can be either the reporter intensity of one or multiple reference channels, or the sum over all sample channels, by selecting all of them as reference channels. in an n-plex in one PSM. The weights wi are the product of the precursor ion intensity exactly at the retention time at which the MS/MS has been recorded times the fill time of the MS/MS spectrum. This is supposed to be proportional to the number of ions that are used for fragmentation. The weights are then exponentiated with a constant which can be set by the user (the ‘isobaric weight exponent’ parameter in the graphical user interface). All calculations are done for raw intensities as well as for intensities corrected for impurities, which are both reported in the output tables.

### Generic data processing

All Andromeda^26^ searches were performed with oxidation of methionine and protein N-terminal acetylation as variable modifications and cysteine carbamidomethylation as fixed modification. Phosphorylation of serine, threonine and tyrosine was additionally added as variable modification for dataset 5. Trypsin was selected as protease allowing for up to two missed cleavages, and the peptide mass was limited to a maximum of 4600 Da. The initial mass tolerance was 20 ppm for precursor ions and 20 ppm for fragment ions. PSM and protein false discovery rates were both applied at 1%. In general, values of parameters in MaxQuant have not been changed from their default values unless explicitly stated. Perseus^27,28^ version 2.1.2 was used for the downstream data analysis, except for the calculation of Wilks’ lambda and the associated p-value, which was done using the R package rrcov: Scalable Robust Estimators with High Breakdown Point^29^ under R version 4.3.2. The graphical abstract is created in BioRender.com. We have uploaded all results to Mendeley Data under doi:10.17632/s3gfmcbghm.1.

### Software development, maintenance and availability

MaxQuant is written in C# under .NET 8. The command line version runs on Windows, Linux and macOS (as long as the vendor libraries for accessing the raw data are compatible). The graphical user interface (GUI) is currently restricted to Windows. Linux and macOS GUI support will be released soon. MaxQuant and Perseus can be downloaded from https://www.maxquant.org/ and are freeware which can be used without restrictions, in particular also for commercial purposes. The code is partially open source (https://github.com/JurgenCox/mqtools7). Bugs should be reported at https://github.com/cox-labs/CoxLab_Bug_Reporting. The Google groups https://groups.google.com/g/maxquant-list and https://groups.google.com/g/perseus-list serve as discussion forums. Many instructional videos are available on YouTube at https://www.youtube.com/@MaxQuantChannel.

## Results and discussion

### Impurity correction for TMTpro

In MaxQuant isotopic labels can be freely defined while standard TMT and iTRAQ labeling sets are pre-configured and set by single clicks. Impurity correction factors can be specified either by direct editing in the graphical user interface or by import from a template file (Figure 1). Dataset 1 was reprocessed with MaxQuant and Proteome Discoverer (PD) results were obtained from reference^7^ for comparison. We compared the number of human protein groups identified under different carrier levels with MaxQuant and PD (Figure 2A). As the carrier level increases, more protein groups were identified from single cell channels for both software platforms, with MaxQuant consistently identifying more protein groups. Given that slight differences exist between the protein group assembly algorithms in MaxQuant and PD, the conclusion is that number of identifications are similar. The higher carrier level also increased the TMT channel leakage effect^30^ especially on neighboring channels (127N), despite the low proportion of impurities, leading to a significant overestimation of the intensities of the single cell channels (Figure 2B). However, with the impurity correction factors of the TMTpro 16-plex reagents provided by the manufacturer, especially for high carrier levels, MaxQuant significantly reduced the channel leakage effects from the carrier channel, similarly to Proteome Discoverer.

**Figure 1.**
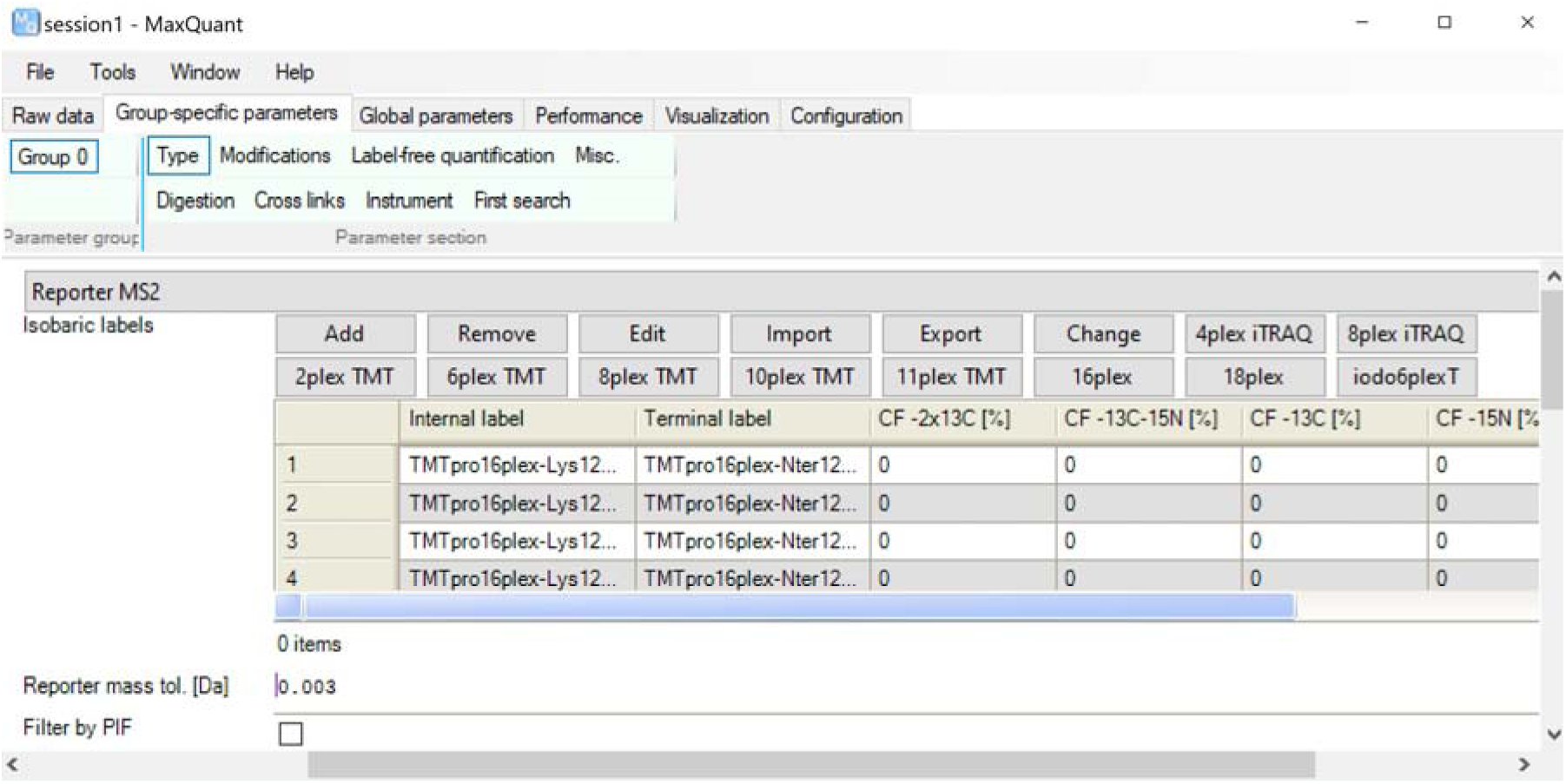
Definition of isobaric labels and impurity correction factors in the graphical user interface of MaxQuant. Clicking on any of the iTRAQ and TMT buttons fills the table with corresponding N-terminal and amino acid-specific labels. Then, impurity correction factors can be defined either by direct editing in the table or by importing from a template file.

**Figure 2.**
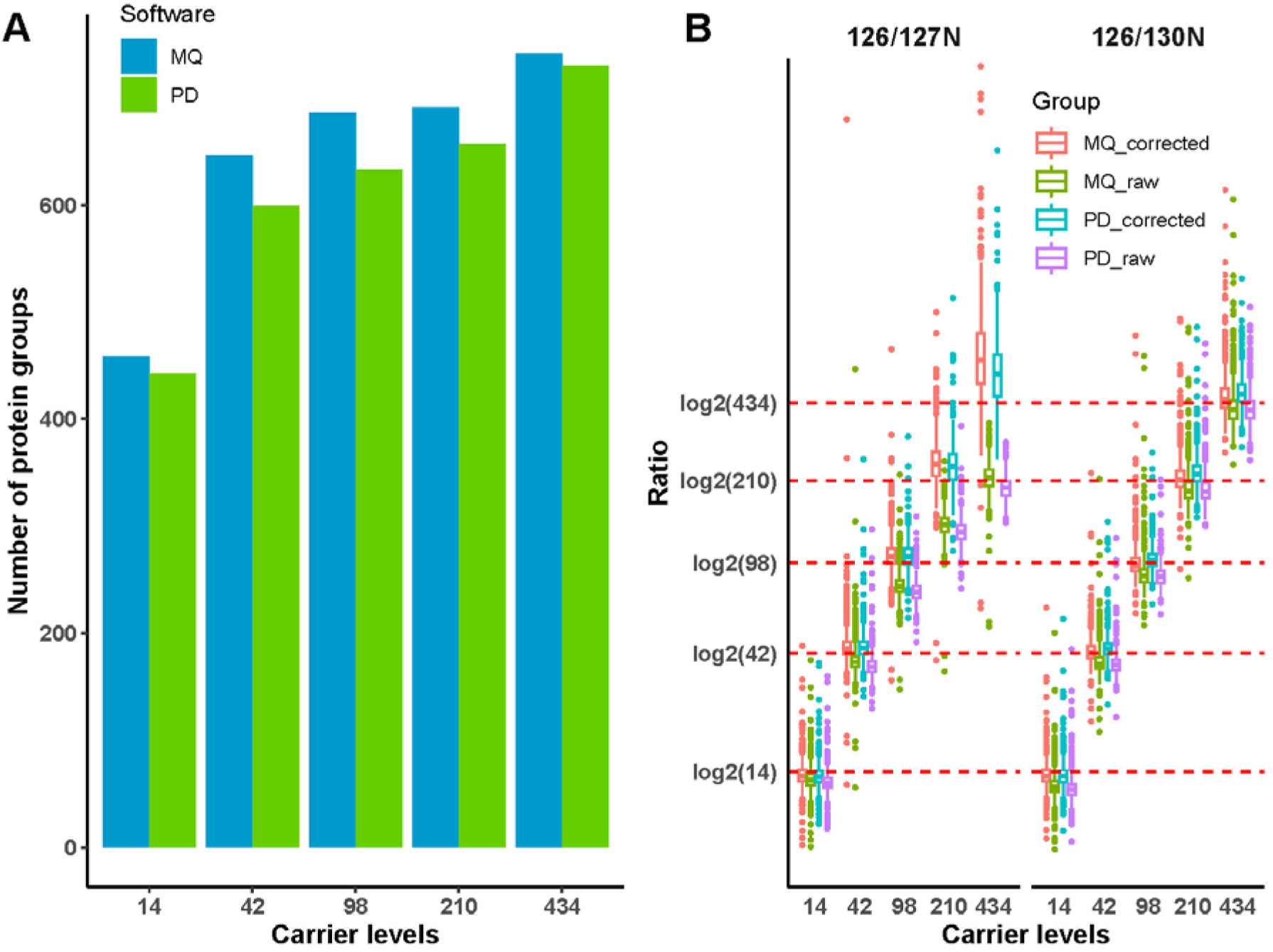
Protein-level identification and quantification results for dataset 1 comparing MaxQuant with the Proteome Discoverer results reported in Ye et al.^7^ **A.** Number of identified protein groups when using human carrier proteome for different carrier levels. Blue: MaxQuant, green: Proteome Discoverer. **B.** Distribution of log2(126/130 N) and log2(126/127 N) for human carrier proteome with and without impurity correction, each with MaxQuant and Proteome Discoverer.

### WM normalization applied to a human body map

In dataset 2, a human organ map of healthy donors is assembled from TMT 10-plex data. We want to apply dimension reduction tools to the high-dimensional proteomics data for data exploration, visualization and hypothesis generation. Figure 3A shows the result of a t-SNE^31^ analysis of the log-transformed protein group reporter ion intensities without applying a normalization method. Reporter intensities from multiple fractions were summed up by defining ‘Experiment’ names accordingly in MaxQuant. Further analysis of the protein group reporter intensities was performed with the Perseus software^27^ including t-SNE analysis through an R-plugin^28^. There are 420 data points in Figure 3A corresponding to 32 organs each from 14 donors. Accordingly, every data point is uniquely assigned to one of the tissues, 10-plexes and donors, the 10-plexes being randomized with respect to the other groupings. Data points are colored by TMT 10-plex in Figure 3A while in Figure 3B the same data is shown, but the coloring is by tissue. Visual inspection indicates that the data clusters by 10-plex and not by tissue. To quantify the clustering preference, we perform multivariate analysis of variance^32^ (MANOVA) on the two-dimensional t-SNE results. We calculate Wilks’ lambda and the associated p-value, once with the 10-plex and once with the tissue as grouping, resulting in Λ = 0.27145, p < 2.2e-16 for 10-plex and Λ = 0.85927, p = 0.513 for tissue, indicating that the batch effect is dominating over the separation by tissue in the unnormalized data. Next, we re-analyzed the data in MaxQuant now using the sum of the two common reference channels for forming ratios to samples, applying WM normalization (Figure 3C,D). The same MANOVA analysis now results in Λ = 0.84639, p = 0.1044 for 10-plex and Λ = 0.030942, p = < 2.2e-16 for tissue. These calculations confirm the visual impression that WM normalization using ratios to the reference channel strongly reduced the 10-plex batch effect and allows partial clustering by tissue. Finally, we repeat the MaxQuant analysis also with WM normalization, but this time ignoring the reference channels and instead taking ratios of samples to the sum of all samples in each 10-plex (Figure 3E,F). This resulted in Λ = 0.88986, p = 0.734 for 10-plex and Λ = 0.0021157, p = < 2.2e-16 for tissue, indicating that the TMT batch effect is mostly removed and that the arrangement of data points in the t-SNE is mostly determined by the tissues, in particular better than when the reference channel was used. Hence, the dominance of biological effect over batch effect is much more pronounced in the analysis that ignored the reference channels, suggesting that the inclusion of these in the experimental design was unnecessary for this dataset. Comparing Figure 3F to the t-SNE map obtained in reference^13^ we conserve similar neighborhood relationships between physiologically related tissues, such as skeletal muscles, heart and arteries from various body parts.

**Figure 3.**
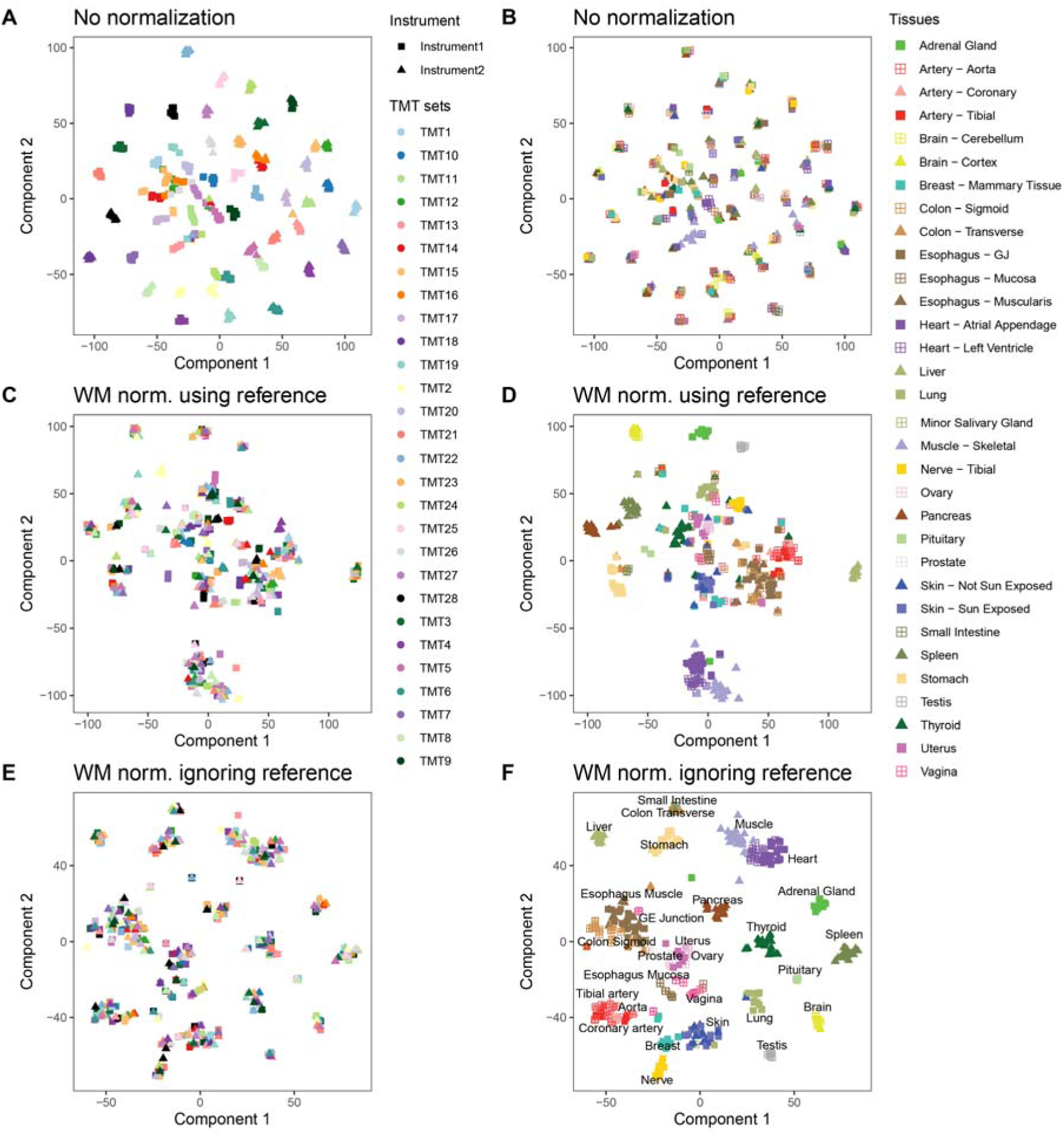
t-SNE analysis of the protein group reporter intensities of dataset 2 Each data point corresponds to a sample. Fractions from the pre-fractionation are summed in MaxQuant. Reference samples are not shown. **A.** t-SNE results when no normalization is applied to the data. Data points are colored according to the TMT 10plex they belong to. **B.** Same as in panel A with coloring according to tissue of origin. **C.** Same as panel A but with WM normalization using the ratio to the reference sample. The coloring refers to 10plex. **D.** Same as in panel C with coloring according to tissue of origin. **E.** Same as panel A but with WM normalization using the ratio to the sum of all tissue samples. The coloring refers to 10plex. **F.** Same as in panel E with coloring according to tissue of origin.

To further investigate the impact of different normalization methods on measurements of individual proteins, we examined four key proteins from the branched-chain amino acid (BCAA) pathway: BCAT1, BCAT2, BCKDHB, and PPM1K, which were also examined by Jiang et al.^33^ The relative abundance of these proteins in different tissues is presented in Figure 4(A-C). The BCAT2 displayed high tissue specificity in adrenal gland, heart ventricle, pancreas and stomach in WM normalized data with (Figure 4B) or without reference channels (Figure 4C), whereas the tissue specificity was not shown in unnormalized data (Figure 4A). All four tissues are among the top five in which BCAT2 is highly expressed, according to the Human Protein Atlas^34^ (Figure 4D). The remaining choroid plexus, which ranks top, was not included in our dataset.

**Figure 4.**
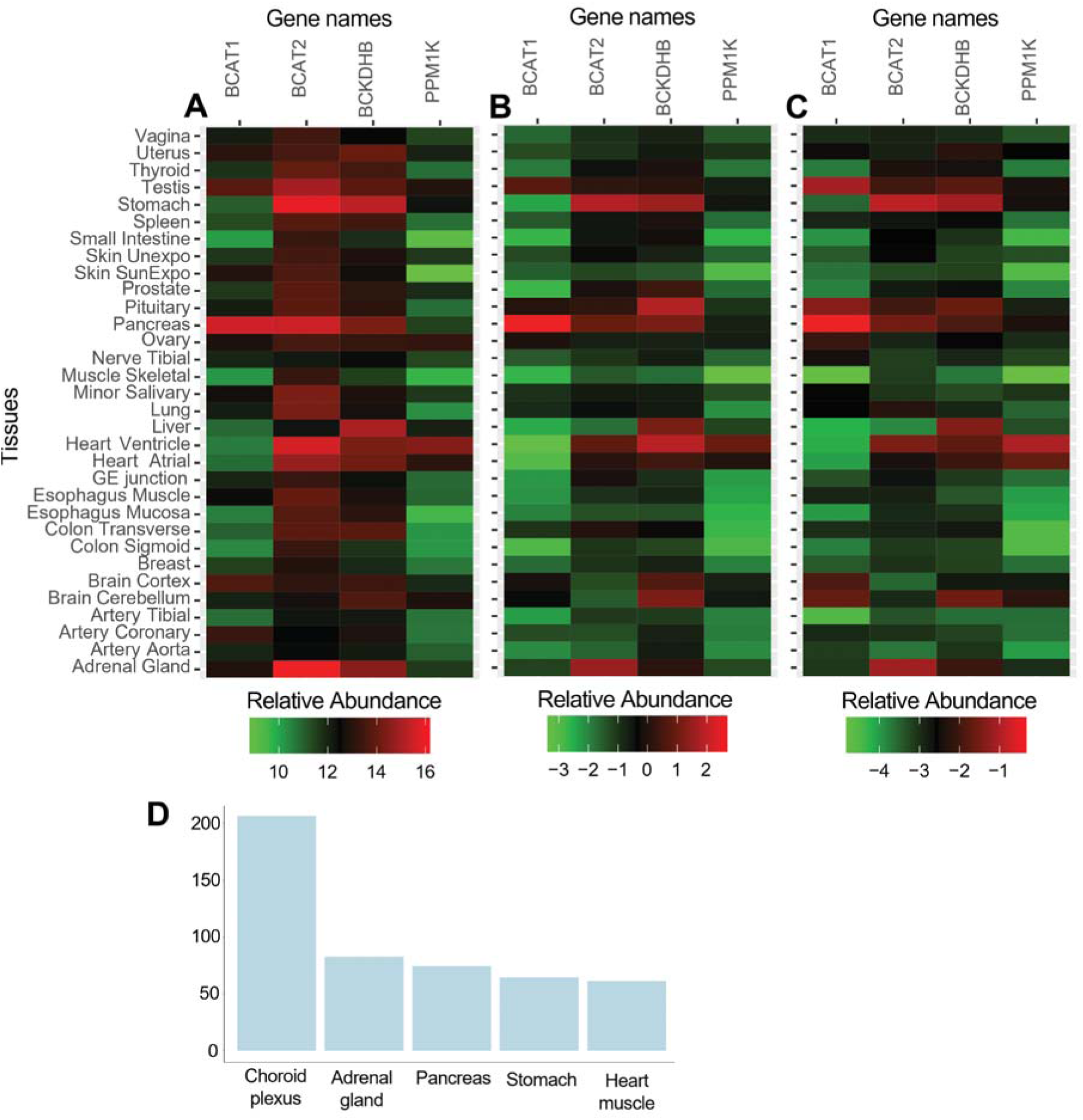
Analysis of key proteins in BCAA pathway. **A.** Heatmap of the relative abundance when no normalization is applied. **B.** Heatmap as A but with WM normalization using the ratio to the reference samples. **C.** Heatmap as A but with WM normalization using the ratio to the sum of all tissue samples. **D.** Barplot of top five tissues in which BCAT2 is highly expressed. nTPM: normalized protein-coding transcripts per million.

While we have used t-SNE so far, several alternative methods exist for dimension reduction, such as principal component analysis (PCA) and Uniform Manifold Approximation and Projection^35^ (UMAP). Comparative studies of these and other techniques exist for transcriptomic data^36^ and cytometry by time-of-flight data^37^. It is not^34^ a priori clear which method performs best on a given proteomics dataset. Hence, we repeated the analysis with PCA and UMAP (Figure 5). Principal component analysis does not result visually in the formation of tissue clusters (Figure 5A). This is due to the fact that a linear projection is not sufficient for reducing the tissue protein expression into a two-dimensional representation. Figure 5B,C show UMAP and t-SNE results, which both find a nonlinear mapping into a 2-dimensional representation. For both methods a clustering by tissue is observed. However, the representation generated by t-SNE is more appealing for interpretation, since the tissue clusters are more evenly spread out over the plot.

**Figure 5.**
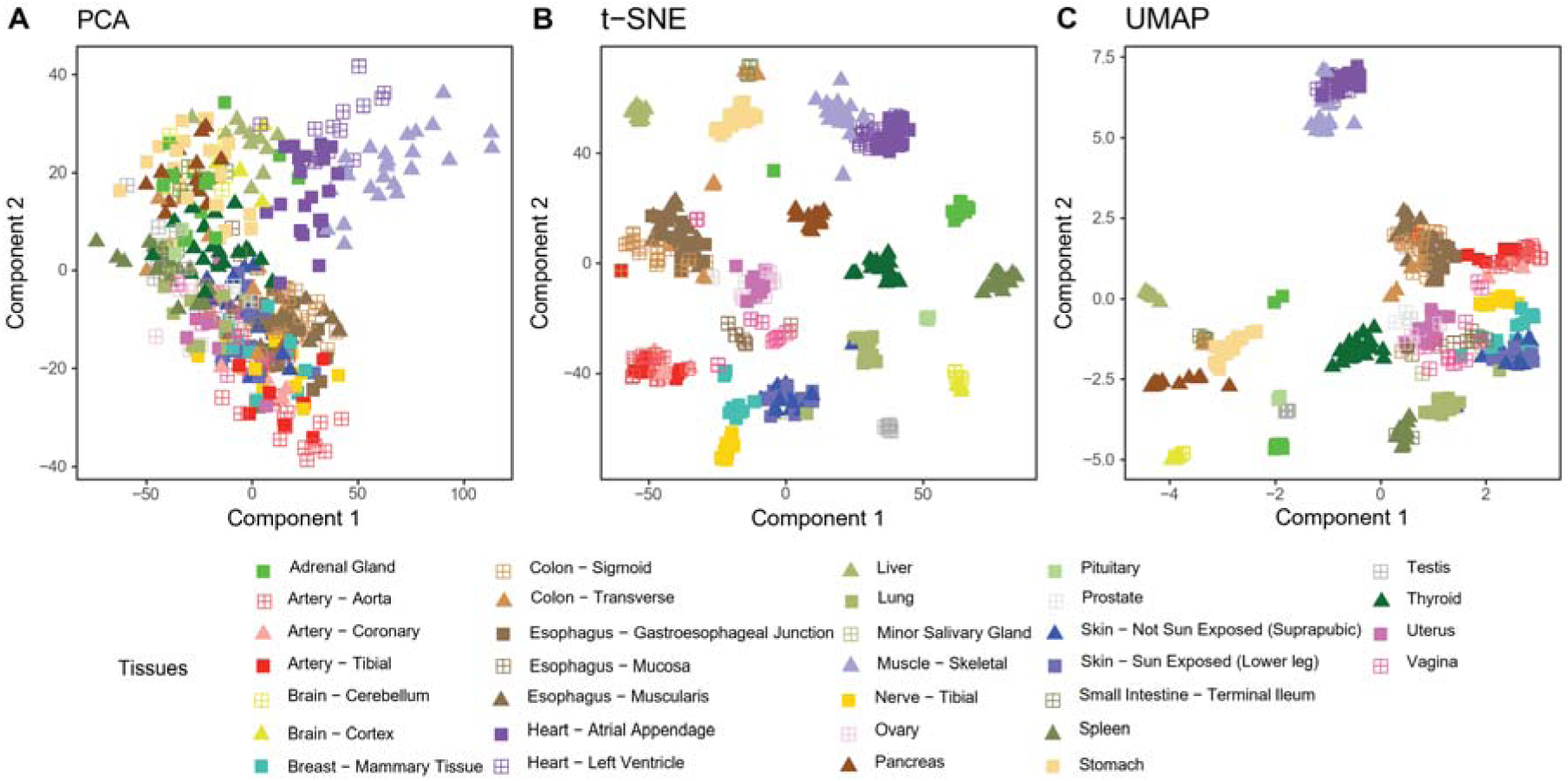
Comparison of dimension reduction methods. **A.** Same data as in Figure 3 but analyzed with PCA instead of t-SNE. **B.** t-SNE data from Figure 3f copied for ease of comparison. **C.** Same data with UMAP.

### WM normalization applied to a two-proteome benchmark dataset

Next, we re-analyze dataset 3, a HeLa-yeast mixture benchmark dataset with several known relative concentrations to study the effect of WM normalization. As for the human tissue map, we want to make use of dimension reduction to study the effects of normalization. While PCA was not able to capture the main effects in the tissue data, due to its linearity, here we expect that PCA is sufficient, since the number of data points (samples) is much lower. Figure 6 shows the first two principal components for several combinations of normalizations, 11-plex or 16-plex and MS2 or MS3 level quantification. In the first row of plots, no normalization is applied and the data is strongly dominated by the n-plex the sample belongs to for all combinations of 11-plex/16-plex and MS2/MS3. In particular, the sample groups are not separated in the first principal component. As a measure for how well the first principal component separates the sample grouping, we calculate -log10 of the ANOVA p-value of the sample grouping in this first principal component (horizontal axes in Figure 6). Without normalization these values range from 0 to 0.05 confirming that there is no significant separation of sample groupings due to the dominance of TMT plex grouping. The middle row of plots in Figure 6 corresponds to applying WM normalization making use of the reference channel. Visually the sample groups are well separated in the first component, while the second component still retains some of the TMT plex grouping. The first component - log10 ANOVA p-values now range from 21.02 to 39.13 confirming significant separation. In the lowest row of Figure 6 we performed WM normalization ignoring the reference channel and using the sum of all sample channels within each TMT plex instead. The -log10 ANOVA p-values now range from 35.88 to 50.10. In all four direct comparisons the significance is higher when the reference channel is not used. This indicates that also in this dataset the inclusion of a reference channel was unnecessary when using WM normalization in MaxQuant.

**Figure 6.**
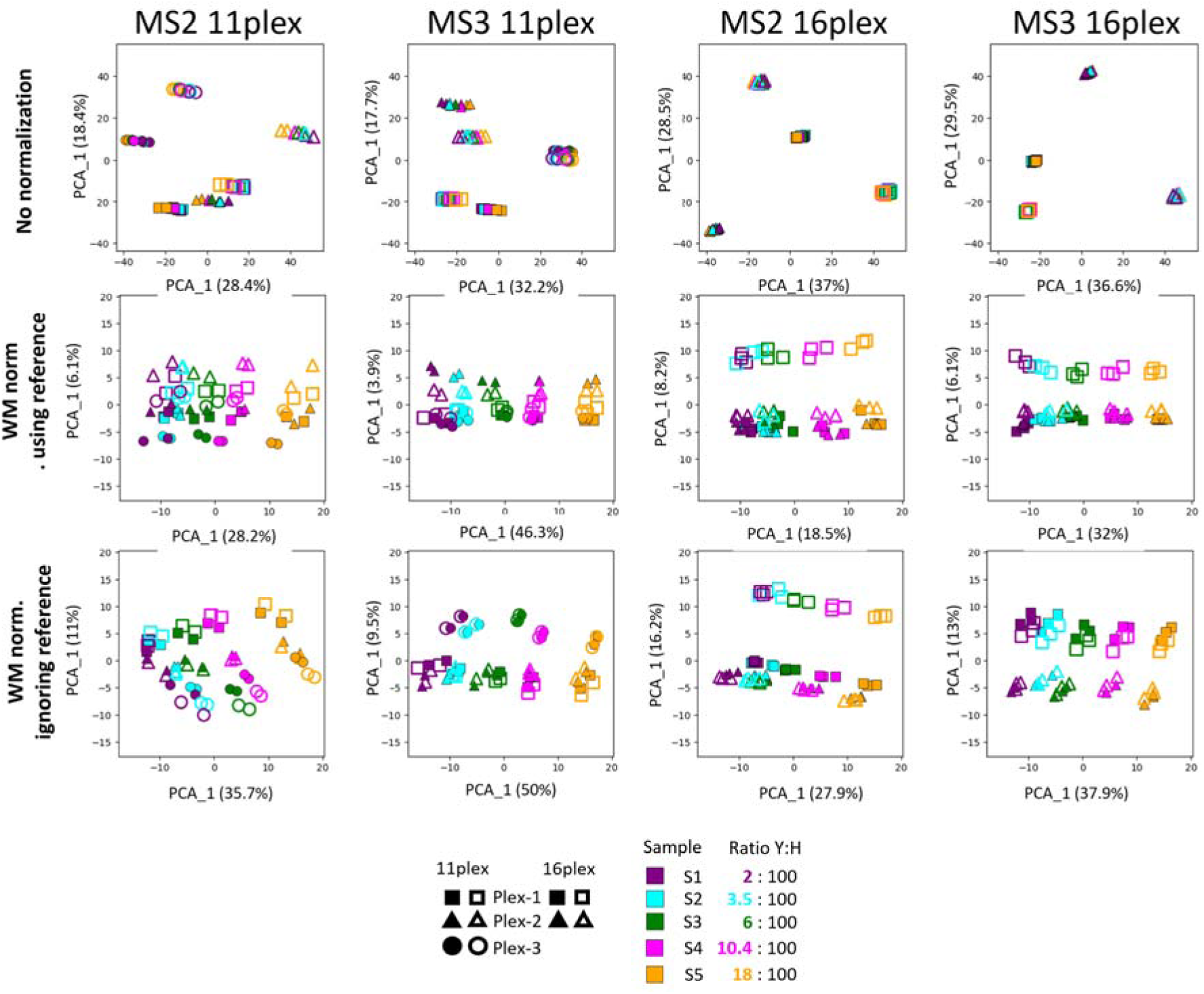
PCA analysis of protein group reporter intensities of dataset 3 for different combinations of normalization, multiplex sizes and MS levels. In the first row, no normalization was applied, in the second row WM normalization using the reference channel and in the last row WM normalization ignoring the reference channel. The number in each plot is -log10 of the ANOVA p-value for the separation of the sample grouping by the first principal component (horizontal axes).

### TMT combined with FAIMS

To demonstrate the feasibility of analyzing FAIMS data combined with TMT labeling we processed dataset 4, a comparative study between MS3 SPS and MS2 FAIMS based TMT quantification. PEAKS (version 8.5) results were taken from the supplementary material of reference^19^. Figure 7A shows the number of peptide identifications for the different technologies and injections, with similar values observed for MaxQuant compared to PEAKS. The same is the case for the number of protein group identifications (Figure 7B). In Figure 7C we show the log2 fold changes in the 9h timepoint between SPS and FAIMS using MaxQuant and the same for PEAKS in Figure 7D, indicating that quantitative results are similar between the software platforms.

**Figure 7.**
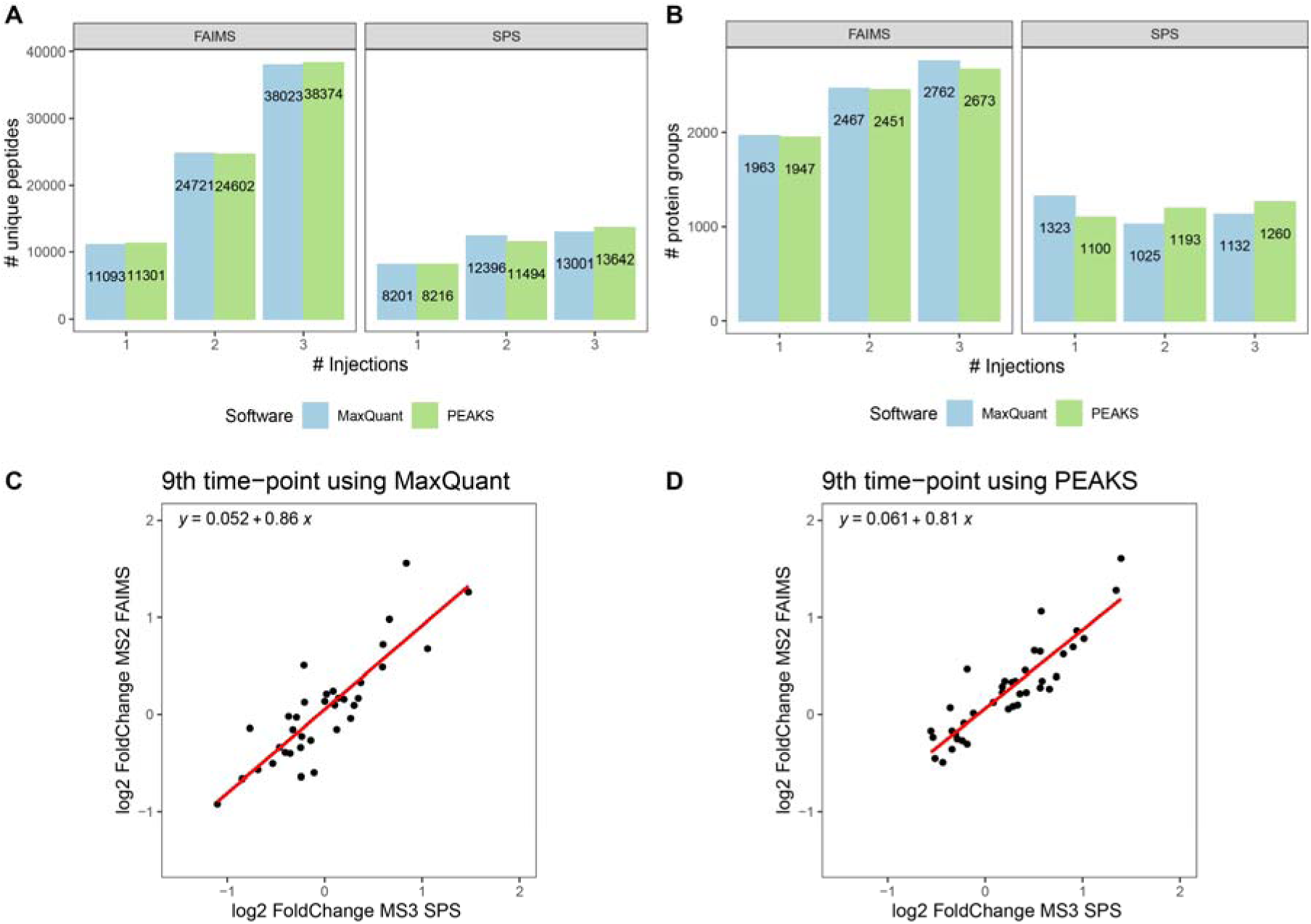
Comparison of MaxQuant and PEAKS applied to dataset 4. **A.** Number of identified peptides for SPS and FAIMS. **B.** Number of identified protein groups for SPS and FAIMS. **C.** Log2 fold change to the 9^th^ time point compared between SPS (horizontal axis) and FAIMS (vertical axis) for MaxQuant. **D.** Same as in C, obtained with PEAKS.

### Analysis of posttranslational modifications

As an example for the analysis of posttranslational modifications (PTMs) with isobaric labels in MaxQuant we selected dataset 5, which is a phosphoproteomics study performed with TMTpro 16-plex. It is a two-species mixture with several known relative concentrations, which is suitable for systematically studying ratio compression. There are two tables in the MaxQuant output that contain the main results regarding identification and quantification of PTMs: the site table, e.g. phospho(STY)Sites.txt, and the modificationSpecificPeptides.txt file. In the former table, each row corresponds to a unique phosphorylation site on a protein and in the latter each row is a peptide with a specific number of modifications. Both tables contain reporter ion intensities across multiple n-plexes if applicable and are suitable for being uploaded to Perseus for multivariate analysis. As an exemplary data analysis we study the effect of applying precursor intensity fraction^38^ (PIF) filtering. MS/MS spectra are filtered for quantification based on the fraction of precursor ions in the isolation window that originate from the peptide that was intended to be fragmented. Figure 8 shows the difference of medians of yeast and human populations for three expected ratios as a function of the PIF threshold that was applied. It can be seen that as the PIF threshold approaches 100% and becomes more stringent, the ratios almost reach their expected values if there was no ratio

**Figure 8.**
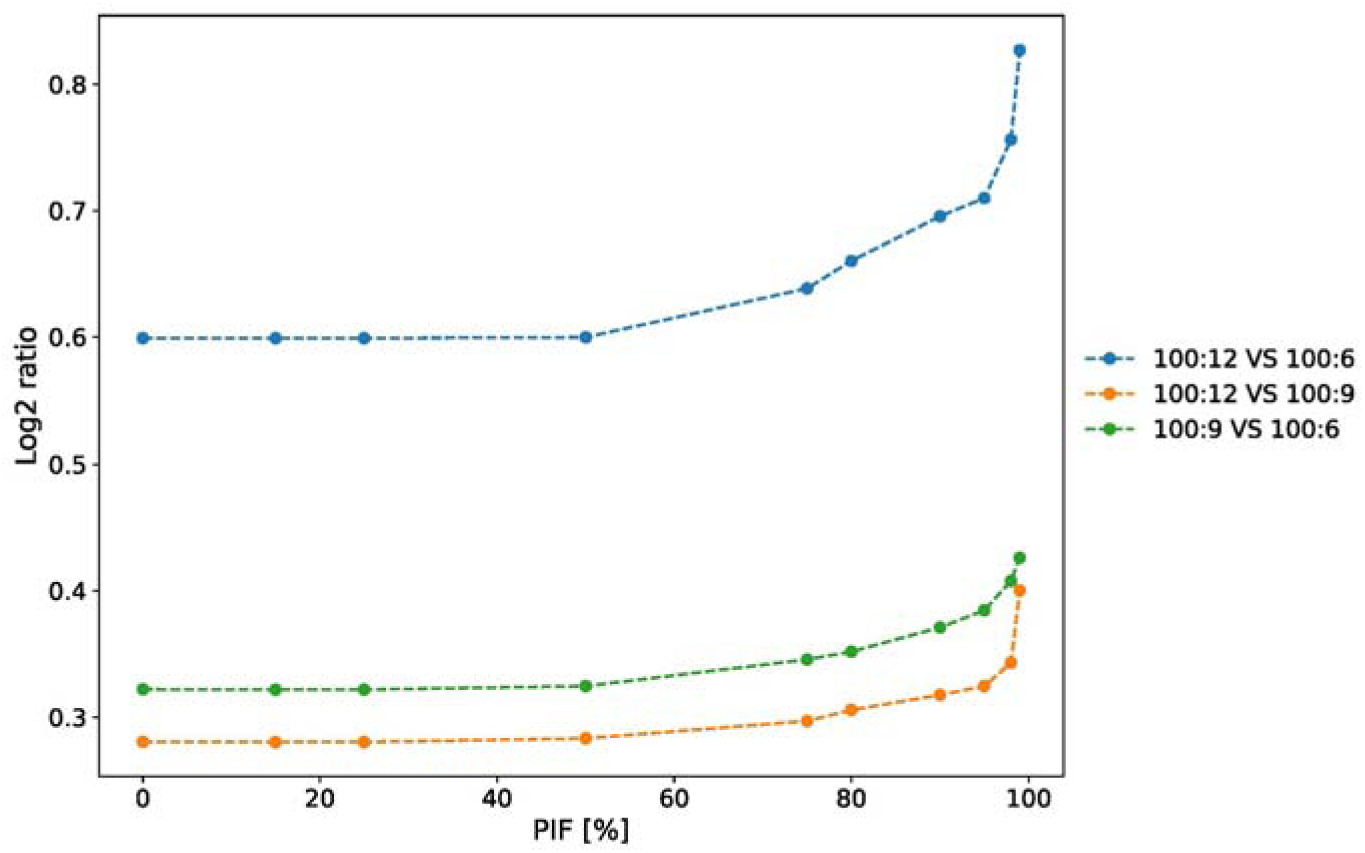
Difference of medians of yeast and human log2 reporter ion intensities for three expected ratios as a function of the PIF threshold that was applied. compression.

## Conclusions

We have shown that the isobaric labeling workflow in MaxQuant performs well on a diverse collection of datasets in terms of identification and quantification. To limit the number of aspects that needed to be compared, we did not use isobaric matching between runs^6^ throughout the manuscript. When activated, it will increase the number of quantified n-plexes and reduce the fraction of missing values. Further increase in identification is expected by integrating spectral prediction^39,40^ with TMT labeling-specific models into the Andromeda scoring in upcoming versions. Alternatively, one can boost identifications with large pre-trained models for peptide-spectrum matching^41^. The development of higher sample plexing reagents would not only reduce analysis time, but also increase the depth of proteome coverage, resulting in a higher number of quantified proteins^9^. In future releases, MaxQuant will also support higher sample plexing, such as TMTpro 32- and 35-plex^42^.

In the datasets under consideration we found that using WM normalization without the reference channel(s) resulted in better results than using the ratios of the sample channels to the reference channel signals. This was the case even though in the human body map even two reference channels were used. While this is not a proof that this will be the case for every dataset in the world, it is a strong indication that reference channels may not be necessary when proper data processing is applied. This frees up more channels for the actual samples and avoids the problem of potential absence or low abundance of proteins in the reference due to the dilution by mixing all samples for the reference. However, one would expect that the normalization without a reference works only if some preconditions are met. The samples in each individual plex should constitute a fairly representative subset of the entire dataset, which requires proper randomization of the experimental design on the plexes. Furthermore, the number of samples per plex should have an effect. If one considers the hypothetical case of plexes containing only two samples, the normalization would certainly fail, while with 35-plexes the no-reference approach should work even better.

Regarding the choice of dimension reduction algorithms, PCA is usually preferable and sufficient if the number of samples is small. In that case it is unlikely that algorithms capable of nonlinear reductions will actually find meaningful nonlinearities due to the lack of data points. When there are sufficiently many samples, as for the human tissue map, nonlinear methods are superior to PCA. Between UMAP and t-SNE, for our particular case here, we found that the t-SNE map was more straightforward to interpret, since it resulted in a more clear-cut visual clustering into tissue groups compared to UMAP. From our experience with other proteomics datasets, this is not always the case and it is best to try both and compare. This is even more recommended, since the differences in results between t-SNE and UMAP are partially due to the different choices of initialization of data points in different implementations^43^.

## Acknowledgements

This work was supported by the German Ministry for Science and Education funding action CLINSPECT-M [FKZ 161L0214E]. The authors thank Michael Bremang and Zilu Ye for their valuable input on the experimental design of the re-analyzed datasets. S.U. and C.D.N. are supported by the Marie Skłodowska Curie European Training Network ’PROTrEIN’ (grant agreement No. 956148). P.K. is supported by the Marie Skłodowska Curie European Training Network ‘PUSHH’ (grant agreement No. 861389).

The authors have declared no conflict of interest.

